# go3: A Fast and Lightweight Library for Semantic Similarity of GO Terms and Genes

**DOI:** 10.1101/2025.09.04.669468

**Authors:** Jose L. Mellina-Andreu, Alejandro Cisterna-García, Juan A. Botía

## Abstract

**Motivation:** Calculation of semantic similarity of Gene Ontology (GO) term subsets is a fundamental task in functional genomics, comparative studies, and biomedical data integration. Existing tools, primarily in Python or R, often face severe limitations in performance when scaling to large annotation datasets.

**Results:** We present go3, the first high-performance, Python-compatible library written in Rust that supports multiple semantic similarity metrics for GO terms and genes. go3 supports both pairwise and batch computations, optimized using Rust’s parallelism and memory safety. Compared to GOATOOLS, the state of the art, it achieves up to 5x speedup and 25x lower memory footprint when loading the GO ontology and gene annotations, and up to x 10 ^3^ speedup when calculating semantic similarities between genes, while preserving output compatibility.

**Availability and implementation:** go3 is implemented in Rust and available through Python 3. It is accessible in GitHub: (https://github.com/Mellandd/GO3).

## Introduction

Gene Ontology (GO) [1], [2] is one of the most widely adopted structured vocabularies in biology, used for representing molecular functions, biological processes, and cellular components of gene products. To quantify the functional similarity between gene products, numerous semantic similarity metrics have been proposed, many relying on GO’s hierarchical structure and annotation corpus [3].

Metrics like Resnik [4], Lin [5], Jiang-Conrath [6], and Wang [7] are frequently used to assess GO term similarity, with applications in gene clustering, function prediction, and disease gene prioritization. While Resnik, Lin, and Jiang-Conrath rely on information content to quantify semantic similarity in the GO hierarchy, they differ in how they normalize or combine this information, leading to distinct sensitivity to term specificity. In contrast, Wang’s method is graph-based, weighting the contributions of all ancestor terms. This diversity reflects that similarity in GO can be modeled either as shared information between terms or as structural relatedness of these elements across the term hierarchy, hence the coexistence of multiple approaches. Notwithstanding, most of these approaches require computing the information content (IC), a statistical measure of specificity that captures how informative a GO term is. The IC of a term *t* is typically defined as:

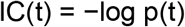

where *p(t)* is the probability of encountering the term *t* in a corpus of annotations, e.g., GAF (Gene Association Files) files. Specific terms deep in the GO hierarchy tend to have higher IC, reflecting more precise biological meaning.

Despite the importance of providing estimates for this functional similarity, for tasks like function prediction or disease gene prioritization, most popular libraries available for this task (e.g., GOATOOLS [8], GOSemSim [9]) are implemented in Python or R, which limits their performance and scalability. Python and R are interpreted and dynamically typed, which introduces runtime overhead in memory allocation, recursion, and data structure traversal, especially when processing large graphs or computing all-vs-all term comparisons.

This limitation becomes critical when researchers aim to compare GO term similarities across multiple gene sets of interest. Such scenarios commonly arise when analyzing statistically significant genes identified in large-scale studies, such as genome-wide association studies (GWAS), to uncover shared biological mechanisms underlying the same phenotype or related disorders. Similarly, transcriptomic analyses often produce numerous gene sets, necessitating millions of GO term comparisons to extract meaningful functional insights. To address this challenge, we developed go3, a semantic similarity engine written in Rust, a systems programming language that offers efficient multithreading. Our library provides a Python interface for easy integration into existing pipelines, combining high-level usability with low-level performance.

### Implementation and features

The go3 library is built entirely in Rust and provides a Python interface through PyO3 bindings, making it both fast and accessible. It implements a suite of well-established semantic similarity measures, including state-of-the-art methods not included in GOATOOLS.

Key features of go3 include:

- Support for multiple semantic similarity measures, including:
  - Node-based: Resnik [4], Lin [5], Jiang-Conrath [6], SimRel [10].
  - Edge and hybrid-based: Wang [7], GraphIC [11], ICCoef [12], TopoICSim [10].
  - Only Resnik, Lin, SimRel and Wang are included in GOATOOLS.
- Efficient GO term and annotation loading, supporting standard OBO and GAF files.
- Automatic IC computation from annotation corpora.
- Batch and parallel computation of term or gene-level similarity using multithreading.
- Memory-efficient internal representations with numeric indexing for GO terms and memoized ancestor caching.

### Benchmark and evaluation

We benchmarked go3 against GOATOOLS [8], which is the established tool for GO analysis in Python. Our benchmarks include ontology and annotation loading time and memory, and pairwise and batch semantic similarity for GO terms and genes. All experiments were conducted on a MacBook Pro M3 Pro (2023) with 18 GB unified memory, macOS 15.5 and python 3.12.2. We used the April 11, 2025 release of the Gene Ontology in OBO format and human annotations from the GOA GAF file. The results obtained were the average of 10 experiments, showing also the confidence intervals on each experiment.

### Ontology and annotation loading

go3 consistently outperformed GOATOOLS in both runtime and memory usage when parsing the GO ontology and GAF annotations, as we can see in Figure 1. This advantage arises from Rust’s compiled execution model and its principle of zero-cost abstractions, whereby high-level constructs (e.g., iterators, traits) are translated into machine code without incurring additional runtime overhead, in contrast to the Global Interpreter Lock (GIL) and dynamic dispatch costs present in Python. Loading times were reduced by up to x5, and memory usage by up to x25, depending on the dataset size.

**Figure 1.**
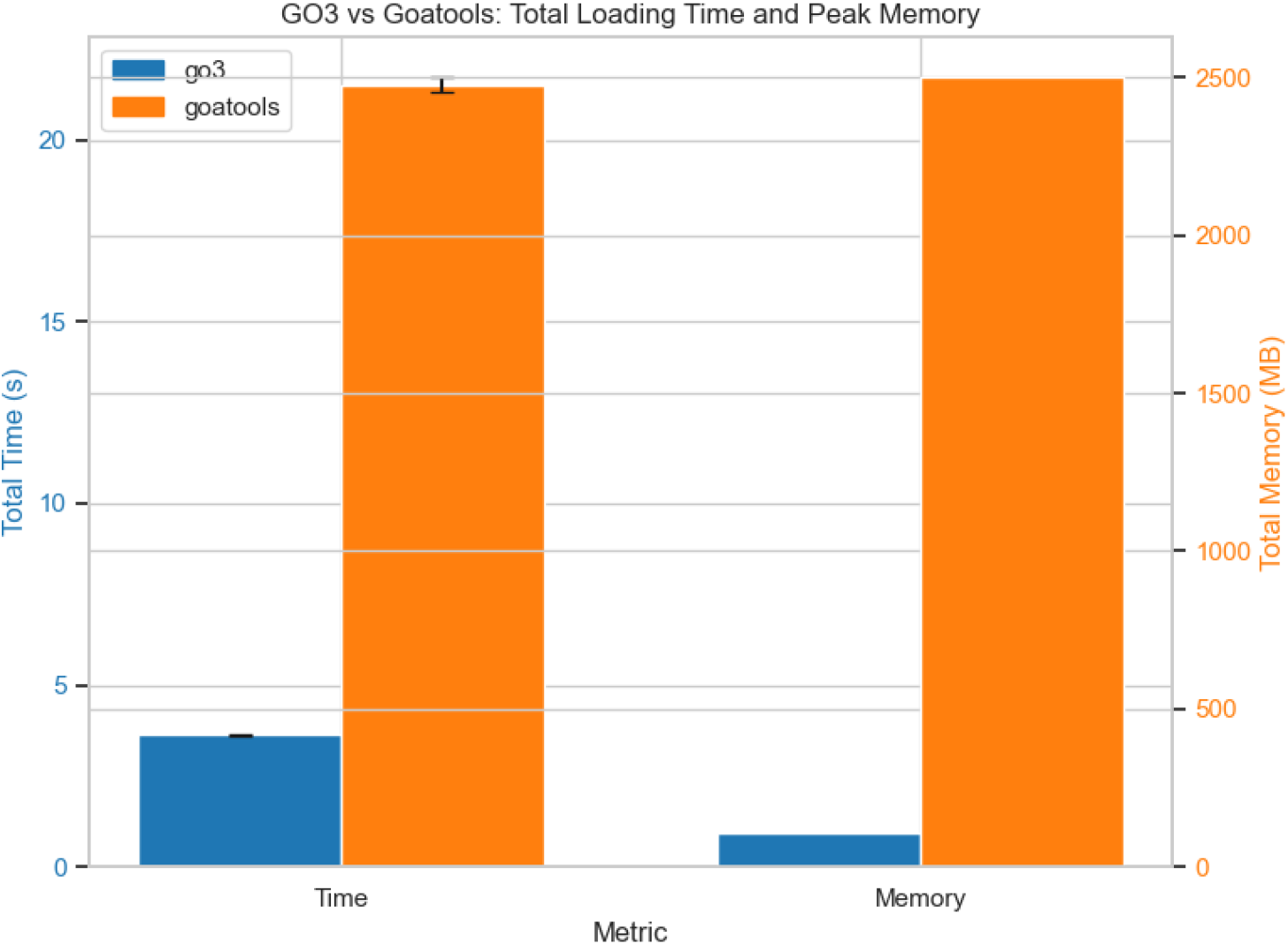
Comparison in loading time and peak memory consumption between GOATOOLS and go3. Average over 10 experiments, showing the confidence intervals at 95% (if it doesn’t appear, then it means that it is too small). Both libraries load the ontology, the annotation (.GAF) files and create the TermCounter element for the IC calculation.

### Semantic similarities

We ran batch semantic similarity computations on random GO term pairs in the BP subontology,the largest in GO, employing several similarity measures implemented in both libraries. In Figure 2. We can see how go3 systematically outperformed GOATOOLS in runtime in all cases, both in term-term similarity and in gene-gene similarity. The difference becomes more noticeable when we calculate semantic similarities between genes, thanks to go3’s built-in functions specific to gene comparisons with default parallelization, generating speedups of up to x 10 ^3^ compared to GOATOOLS. For example, to obtain the semantic distance matrix for 100 genes with Resnik, go3 needs, on average 1. 8 × 10^−4^ s, while GOATOOLS needs 1. 37 × 10^− 1^s.

**Figure 2.**
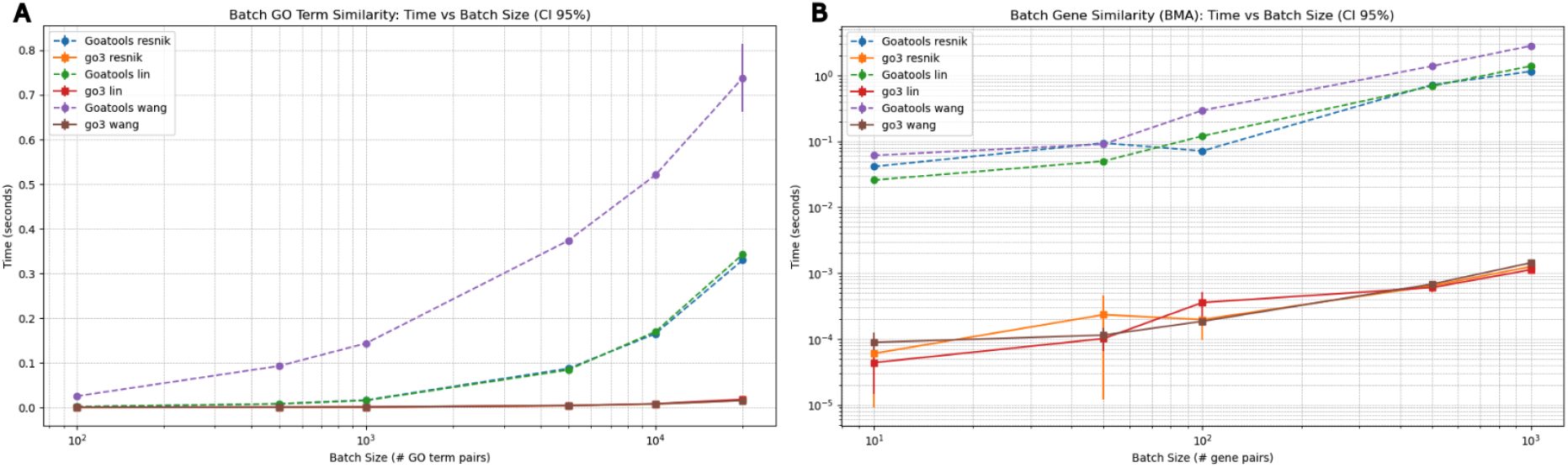
Comparison in runtime between GOATOOLS and go3 when calculating semantic similarities. The results are the average over 10 tries, showing the confidence intervals at 95% (if it doesn’t appear, then it means that it is too small). A) Plot comparing times in term to term similarities, depending on the batch size. B) Plot comparing times in gene-to-gene similarities with bma groupwise aggregation, depending on the batch size.

## Conclusion

**go3** provides a modern, high-performance solution for semantic similarity computation over the Gene Ontology, bridging the gap between usability and scalability. By combining Rust performance with Python usability, it enables efficient large-scale analyses and introduces recent state-of-the-art similarity functions. The library is fully open-source and available via PyPI for seamless integration into computational biology pipelines.

